# Single-nuclei transcriptomics of schizophrenia prefrontal cortex primarily implicates neuronal subtypes

**DOI:** 10.1101/2020.07.29.227355

**Authors:** Benjamin C. Reiner, Richard C. Crist, Lauren M. Stein, Andrew E. Weller, Glenn A. Doyle, Gabriella Arauco-Shapiro, Gustavo Turecki, Thomas N. Ferraro, Matthew R. Hayes, Wade H. Berrettini

## Abstract

Transcriptomic studies of bulk neural tissue homogenates from persons with schizophrenia and controls have identified differentially expressed genes in multiple brain regions. However, the brain’s heterogeneous nature prevents identification of relevant cell types. This study analyzed single-nuclei transcriptomics of ~275,000 nuclei from frozen human postmortem dorsolateral prefrontal cortex samples from males with schizophrenia (n = 12) and controls (n = 14). 4,766 differential expression events were identified in 2,994 unique genes in 16 of 20 transcriptomically-distinct cell populations. ~96% of differentially expressed genes occurred in five neuronal cell types, and differentially expressed genes were enriched for genes associated with schizophrenia and bipolar GWAS loci. Downstream analyses identified cluster-specific enriched gene ontologies, KEGG pathways, and canonical pathways. Additionally, microRNAs and transcription factors with overrepresented neuronal cell type-specific targets were identified. These results expand our knowledge of disrupted gene expression in specific cell types and permit new insight into the pathophysiology of schizophrenia.

## Introduction

Schizophrenia is a chronic psychotic illness affecting ~1% of the population worldwide. Transcriptomic studies utilizing bulk homogenates of frozen human postmortem brain tissue from persons with schizophrenia and controls have identified differentially expressed genes (DEGs) in the amygdala^1^, hippocampus^2, 3^, superior temporal gyrus^4^, anterior cingulate cortex^5, 6^, and dorsolateral prefrontal cortex (dlPFC)^7, 8^, with the largest study identifying ~4,800 DEGs associated with schizophrenia in the dlPFC^9^. However, the heterogenous cellular composition of the bulk homogenates prevents identification of the specific cell types in which relevant genes are differentially regulated and expressed. Two studies used laser capture microdissection followed with transcriptomic analysis by microarray to elucidate the effect of schizophrenia on the transcriptome of individual neural cell types. Examination of layer 3 and 5 pyramidal neurons in the dlPFC identified ~1,400 DEGs^10^ in the context of schizophrenia, whereas a study of parvalbumin positive (PVALB+) interneurons in the dlPFC identified ~900 DEGs^11^. A substantial portion of the DEGs identified in these studies were not detected in previous examinations of bulk homogenates of the same brain regions, suggesting that examination of transcriptomic changes associated with schizophrenia at the level of neural cellular subpopulations is necessary to fully appreciate the neuropathophysiology of the disorder^10, 11^.

Laser capture microdissection studies of human postmortem brain tissue are limited by their ability to examine a small number of cell types in a targeted fashion, relatively low throughput, and the pooling of cells, which loses the variability of the transcriptome between cells and may collapse transcriptomically-distinct subpopulations. Recent advances in single nuclei RNA sequencing (snRNAseq) allow for simultaneous transcriptomic profiling of thousands of nuclei, across all neural cell types in frozen human postmortem brain homogenate, with simultaneous indexing of transcripts at the sample, nucleus, and individual transcript level (unique molecular identifier, UMI). The utility of this approach for human postmortem study is supported by evidence suggesting that single cells and their nuclei have similar transcriptomes, with ~98% of transcripts having the same relative levels^12^. snRNAseq has identified cell type-specific transcriptomic changes in human postmortem brain samples from Alzheimer’s disease^13^, autism^14^, multiple sclerosis^15^, and major depressive disorder^16^.

In this study, we performed snRNAseq of ~275,000 nuclei from dlPFC of individuals with schizophrenia (n = 12) and controls (n = 14). We chose to examine the dlPFC due to the evidence of dlPFC dysfunction in schizophrenia^17^. We identified 4,766 DEGs in 16 of 20 transcriptomically-distinct cell populations. ~96% of the DEGs occurred in five neuronal cell types. The DEGs were enriched for genes associated with schizophrenia GWAS loci and overrepresented in gene ontologies and KEGG pathways previously associated with the pathophysiology of schizophrenia. Canonical pathway analysis identified cluster-specific alterations in metabolic and cell signaling pathways, and microRNA, transcription factor, and upstream regulator analyses identified putative regulators of cluster-specific DEG. Taken together, the results of this study help elucidate the cell type-specific transcriptomic and neurobiological changes that underlie schizophrenia.

## Materials and Methods

### Brain Samples

This study was approved by the University of Pennsylvania Institutional Review Board. Fresh frozen postmortem dlPFC tissue from male individuals with schizophrenia (n = 14) and controls (n =14) were obtained from the Douglas-Bell Canada Brain Bank at McGill University, the Human Brain and Spinal Fluid Resource Center at UCLA and the New South Wales Brain Tissue Resource Center. Schizophrenia cases were individuals who were clinically diagnosed with schizophrenia using DSM-IV criteria and controls were individuals without history of psychiatric disease who died of non-central nervous system-related reasons. All reported age, sex, ethnicity, postmortem interval, and prefrontal cortex pH data are based on associated medical records (Supplementary Table 1). Gray matter samples from the dlPFC were dissected by trained neuroanatomists at their respective brain banks.

### 10x Library Preparation, Sequencing, and Quality Control

Nuclei were isolated from frozen postmortem dlPFC (~30mg) using a modified version of a previously described protocol^16^ (see Supplemental Methods). Microfluidics capture and sequencing library preparation was performed with the 10x Genomics Chromium Single Cell 3’ GEM, Library and Gel Bead Kit v3.0 at the Children’s Hospital of Philadelphia Center for Applied Genomics per manufacturer’s instructions. To achieve a target capture of ~10,000 nuclei per sample, ~20,000 nuclei per sample were loaded. Libraries were sequenced in pools of eight on Illumina NovaSeq 6000 S2 flow cells. Pools contained schizophrenia and control samples, to minimize any batch effects. CellRanger version 3.1 was used to align reads to the hg38 pre-mRNA transcriptome. Filtered read count matrices for all subjects and nuclei were merged into a single Seurat object for subsequent quality control and clustering using Seurat version 3.1. For initial quality control assessment, the distributions of the numbers of genes and UMIs were determined. Nuclei with the lowest 1% of genes (< 470 genes) were removed, as they were unlikely to be informative in downstream analyses. Likewise, nuclei in the top 1% of UMI count (UMI > 60,335) were removed to reduce the presence of multiplets in downstream analyses. Finally, nuclei with >10% of reads from mitochondrial genes were excluded and mitochondrial transcripts were removed from the dataset^18^.

### Calculation of PCs, Clustering, and Cell Type Annotation

Transcript counts were normalized to 10,000 counts per subject and scaled. Variably expressed genes were identified with the FindVariableFeatures function in Seurat using the mean.var.plot selection method and analyzing only genes with mean scaled expression between 0.003 and 2. These parameters identified 2,486 highly variable genes, which were used to generate principal components (PCs). Clustering was performed in Seurat using the first 50 PCs. Initial clustering was performed at a resolution of 0.25. Two schizophrenia samples did not cluster with the other 26 samples and were removed as outliers. The dataset was reclustered and two cell populations with low mean UMIs were removed. Six clusters with >90% of nuclei coming from ≤2 subjects were also removed, with remaining clusters having <30% of nuclei coming from ≤2 subjects. Major cell types and neuronal subtypes were identified using known cell type markers and methods described in Nagy et al.^19^ (see Supplemental Methods). Two clusters with mixed cell type markers were removed for a final total of 20 clusters.

### Differential Gene Expression Analysis

Count data were normalized to one million reads, extracted from Seurat, and converted to log2 counts per million (cpm). Metadata and cpm were merged to form a SingleCellAssay object for each cluster. Genes expressed in <20% of the nuclei in a cluster were excluded from downstream analyses. Differential expression analysis between cases and controls was performed using the *MAST* R package^20^ by fitting the following linear mixed model:

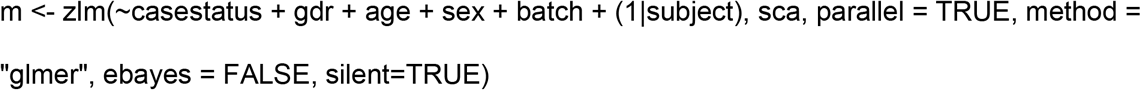

Case status, age, the capture and sequencing batch, and the number of genes detected in each nucleus (gdr) were included as fixed effects. Subject was included as a random effect to account for correlations between the nuclei coming from a single person. To optimize the random and fixed effects coefficients in the penalized iteratively reweighted least squares step, the integer scalar in the *lme4* R package was set equal to zero, as previously described^14^ (https://github.com/DmitryVel/KriegsteinLab/blob/master/snRNAseq_DGE.R). Likelihood Ratio Test was performed in *MAST* to test for differences between the model with and without schizophrenia case status, identifying gene expression differences associated with schizophrenia. DEGs were defined as those that were a) statistically significant after multiple testing correction with false discovery rate (FDR) = 0.1 and b) had at least a 10% difference in expression between case and controls (log_2_ fold change ≥0.14).

### Overrepresentation of GWAS loci

To determine if schizophrenia, bipolar disorder, and Alzheimer’s disease GWAS loci were overrepresented in cluster-specific DEGs, MAGMA ^21^ was used to identify significant genes using GWAS summary statistics. Cluster-specific overrepresentation of GWAS loci was determined by performing a hypergeometric test using the overlapping list of genes that were checked for differential expression in each cluster and were significant by MAGMA analysis.

### Enrichment Analysis with FUMA

Overrepresentation of brain expressed genes was determined by comparing cluster-specific DEGs to brain region-specific transcriptome data from the Genotype-Tissue Expression (GTEx) project (v8)^22^ using FUMA^22, 23^. Similarly, overrepresentation of brain expressed genes throughout the human life span was determined by comparing cluster-specific DEGs to the BrainSpan transcriptomics data ^24^. Identification of gene ontologies, KEGG pathways, transcription factor targets, and microRNA targets that were significantly overrepresented in cluster-specific DEGs was analyzed using FUMA and data from the Molecular and Signatures Database (MsigDB) v7.0 ^25^. For all analyses, all genes, except the MHC region, from Ensembl v92 were used for the gene background and all results were corrected for multiple testing using a Bonferroni correction (α=0.05).

### Canonical pathways and upstream regulators

Canonical pathway and upstream regulator analysis was conducted using Ingenuity Pathway Analysis^26^. DEGs from each of the primary neuronal clusters were analyzed independently using the Core Analysis function and a list of DEG Ensembl IDs, *MAST* raw p-values, FDR corrected p-values (i.e. q-values), and log_2_ fold change. The analyses from the five clusters were compared using the Compare Analyses function. P-values were calculated using a right-tailed Fisher’s exact test and corrected for multiple testing using a Benjamini-Hochberg correction (α=0.1).

### Statistics

Statistical calculations were performed with SPSS (v24).

## Results

### Single nuclei RNA sequencing and identification of cell types

To better understand the effects of schizophrenia on the neural transcriptome, we utilized snRNAseq to profile dlPFC nuclei from frozen human postmortem brain samples from 12 individuals with schizophrenia and 14 controls (Supplementary Table 1). Schizophrenia and control groups did not differ in mean age, postmortem interval, or pH (Supplementary Table 1 and Supplementary Fig. 1). snRNAseq was performed using the 10x Genomics platform and sequenced to an average depth of ~492 million reads per sample. We initially identified 361,681 nuclei, with average medians of ~2,987 genes and ~6,886 UMI per nucleus (Supplementary Table 2). The number of sequencing reads per sample, number of nuclei per sample, average sequencing reads per nuclei per sample, median genes per nuclei per sample, and median UMI per nuclei per sample were consistent between schizophrenia and control groups (Supplementary Fig. 2). RIN value was unavailable for all samples, so RNA quality was evaluated by examining sequencing-derived surrogates of RNA quality from CellRanger. RNA quality did not differ between schizophrenia and control groups, with no difference in the fraction of reads mapped confidently to the genome, intergenic regions, exonic regions, intragenic regions, or the transcriptome (all Mann-Whitney U p-values >0.28; Supplementary Table 3). After quality control measures were applied, we identified 273,050 high confidence nuclei: 145,120 from control samples and 127,930 from schizophrenia samples. These nuclei were used for all further analyses.

To identify nuclei from transcriptomically-distinct cellular populations, we used the snRNAseq counts for nuclei from all individuals to perform unbiased clustering using Seurat^27^, and 20 cellular clusters were identified (Fig. 1a and Supplementary Fig. 3). Clusters were annotated by expression of known cellular subtype markers and all major neural cell types were identified (Fig. 1a–b). The schizophrenia and control groups were consistent for the number of nuclei from major cell types (Fig. 1c) and the number of nuclei per cluster (Fig. 1d & 1e). Excitatory neuron (ExNeuro) and inhibitory neuron (InNeuro) clusters were further annotated using known markers for cortical layers and defined subpopulations (Fig. 1b and Supplemental Fig. 4a-b).

**Fig. 1:**
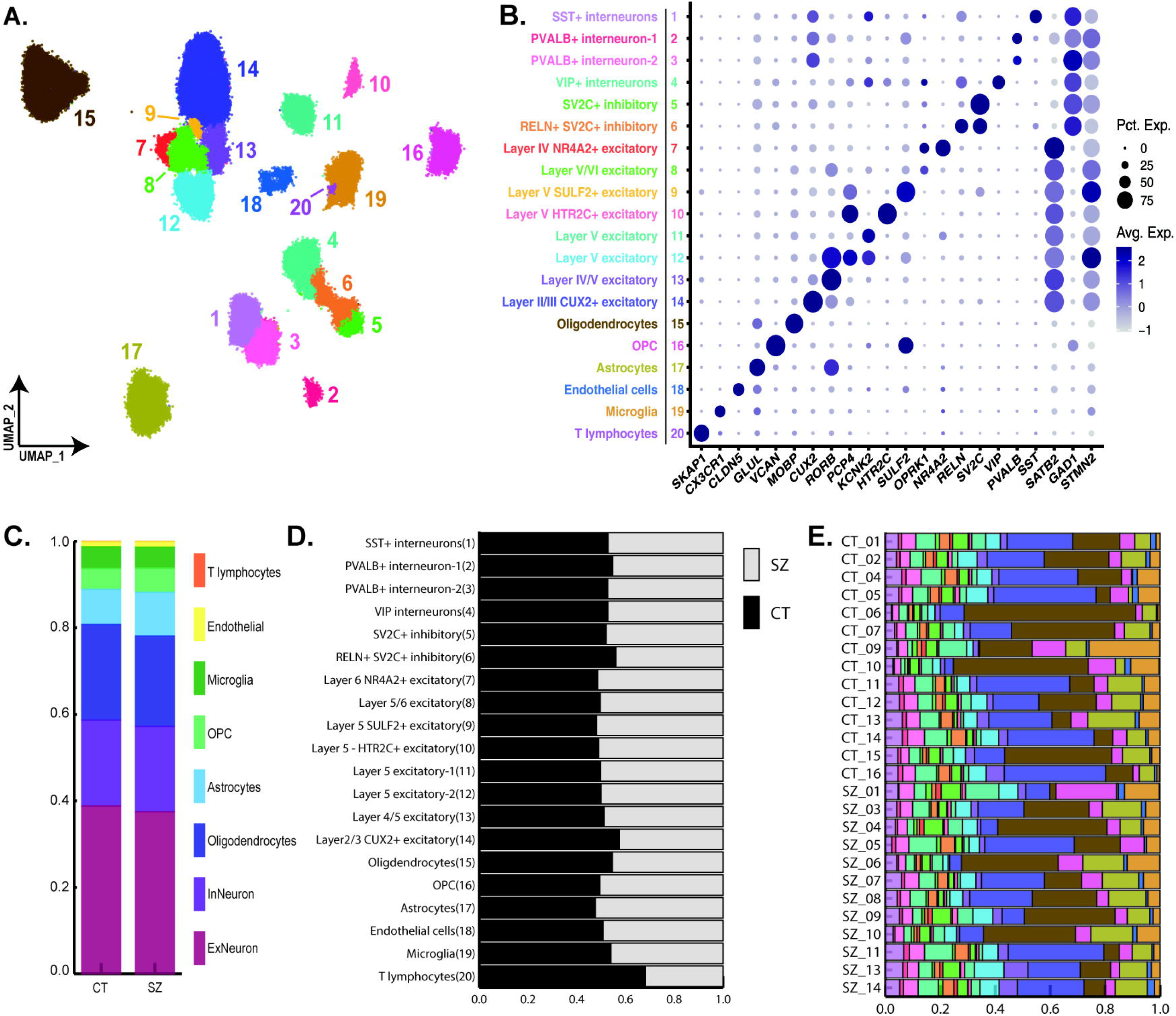
snRNAseq and clustering. **a** The transcriptome profile of ~275,000 nuclei were utilized for unbiased clustering and is presented as a uniform manifold approximation and projection (UMAP) dimension reduction plot of all nuclei color coded by cluster. **b** Clusters were annotated with genes known to be markers for major neural cell types. The size and color of dots is proportional to the percentage of cells expressing the gene (Pct. Exp.) and the average expression level of the gene (Avg. Exp.), respectively. The cluster numbers and colors are matched to that of the UMAP. **c** The proportion of major cell types between the schizophrenia and control groups. **d** The proportion of schizophrenia and control nuclei in each cluster. The labels and numbers correspond to those of the UMAP and dot plot. **e** Cluster contribution by individual sample. Colors correspond to those of the UMAP and dot plot. Abbreviations: somatostatin (SST), parvalbumin (PVALB), vasoactive intestinal peptide (VIP), synaptic vesicle glycoprotein 2C (SV2C), reelin (RELN), nuclear receptor subfamily 4 group A member 2 (NR4A2), sulfatase 2 (SULF2), 5-hydroxytryptamine receptor 2C (HTR2C), cut like homeobox 2 (CUX2), oligodendrocyte precursor cells (OPC).

### Differential expression

Cell type-specific changes in nuclear transcript levels were compared between the schizophrenia and control groups using a hurdle model, a linear mixed model, in MAST^20^. 4,766 differential expression events were identified in 2,994 unique DEGs (q-value <0.1; expression change ≥10% Supplementary Table 4). DEGs were detected in 16 of 20 cellular clusters, with 35.3% of differential expression events being upregulation and 64.7% downregulation (Fig. 2a). The number of DEGs per cluster was unrelated to the number of nuclei per cluster (r(18)=0.16, p=0.50, Pearson’s Correlation; Supplementary Fig. 5). Of the unique DEGs, 96.2% occurred in five neuronal cell types (primary neuronal cell types), including four clusters of ExNeuro (cluster 10 layer 5 HTR2C+ ExNeuro, cluster 12 layer 5 ExNeuro, cluster 13 layer 4/5 ExNeuro, and cluster 14 layer 2/3 ExNeuro) and a PVALB+ InNeuro cluster (cluster 3; Fig. 2b).

**Fig. 2:**
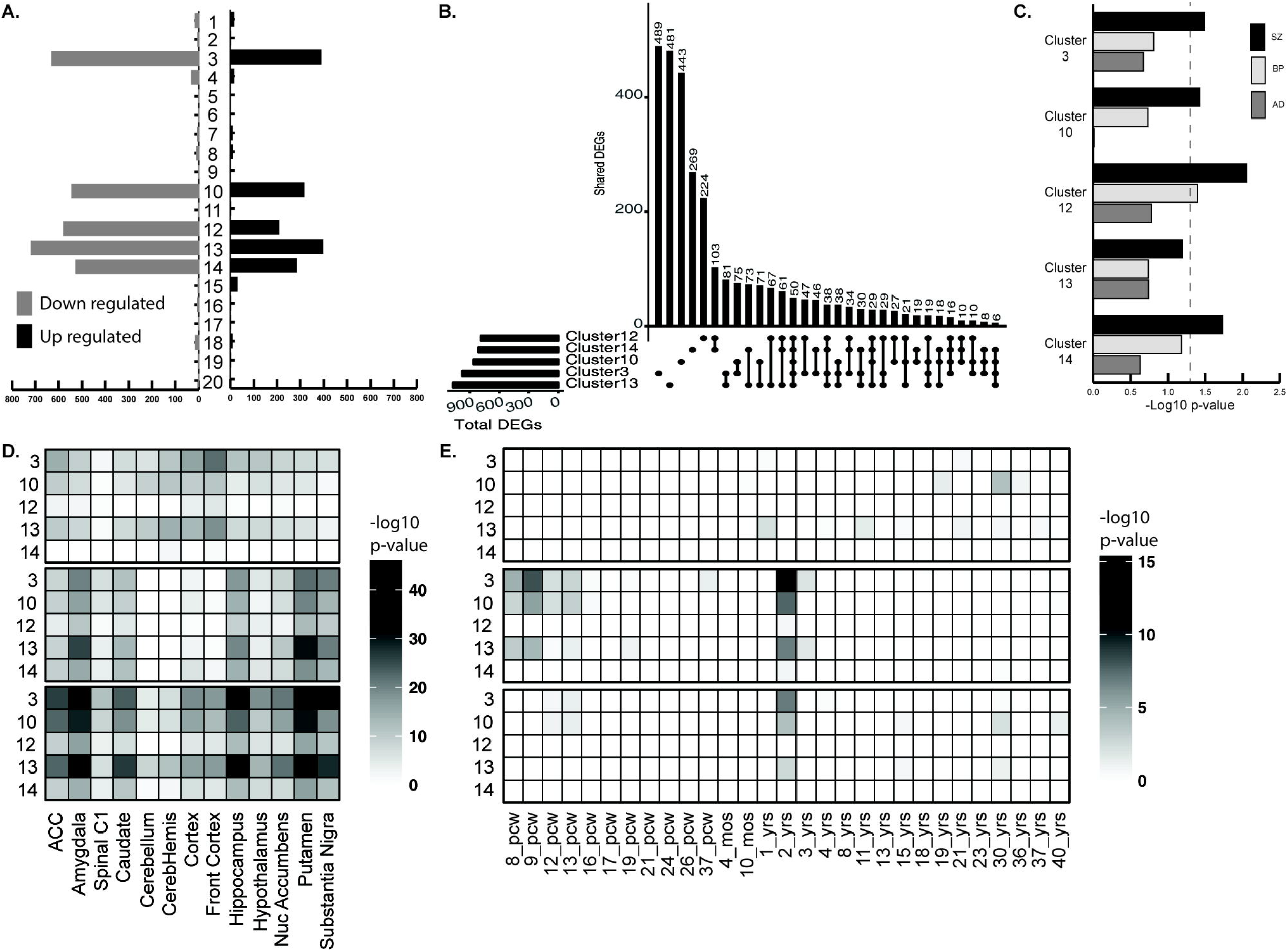
Differential expression. **a** The number of up or down regulated genes by cell type. **b** UpSet plot of the number of unique differentially expressed genes (DEGs) that are unique or shared between the five neuronal cell clusters that encompass the majority (~96%) of all DEGs. **c** Overrepresentation of schizophrenia (SZ), bipolar disorder (BP), and Alzheimer’s disease (AD) GWAS loci in primary neuronal clusters. **d** Heatmap of the overrepresentation of cluster-specific DEGs throughout the brain. **e** Heatmap of the overrepresentation of cluster-specific DEGs throughout the neural life span.

### DEGs associated with GWAS loci

MAGMA was used to determine if cell type-specific DEGs were enriched for genes identified in GWAS of schizophrenia^28^ or bipolar disorder^29^, because previous studies identified substantial heritability overlap^30^. Schizophrenia GWAS loci-associated genes were overrepresented in four of five primary neuronal cell types, with cluster 12 also showing overrepresentation of bipolar disorder GWAS genes (Fig. 2c and Supplementary Table 5). Supporting the specificity of these observations, no enrichment was detected for Alzheimer’s disease GWAS genes (Fig. 2c and Supplementary Table 5). Similarly, cell type-specific DEGs were enriched for GWAS Catalog genes for schizophrenia in cluster 3 PVALB+ InNeuro and cluster 13 layer 4/5 ExNeuro and for bipolar disorder in cluster 10 layer 5 HTR2C+, cluster 12 layer 5, and cluster 13 layer 4/5, and cluster 14 layer 2/3 ExNeuro (Supplementary Table 6).

### Overrepresentation of brain expressed and developmental genes

Comparison of cluster-specific DEGs to brain region-specific transcriptomic data from GTEx shows substantial overrepresentation of up and down regulated DEGs throughout the brain for the primary neuronal clusters (Fig. 2d and Supplementary Table 7). Intriguingly, when compared to BrainSpan transcriptomic data, overrepresentation of cluster-specific DEGs was concentrated at eight to thirteen post-conception weeks and two years of age (Fig. 2e and Supplementary Table 8), suggesting a putative neurodevelopmental relevance.

### Gene Ontologies and KEGG pathways

To understand the neurobiological consequences of the DEGs in the primary neuronal clusters, FUMA was used to identify gene ontologies (GO; Supplementary Tables 9) and KEGG pathways (Supplementary Tables 10) in which DEGs from each cell cluster were over- or under-represented. GO and KEGG analysis identified cluster-specific representation changes of DEGs in ontologies and pathways previously associated with the pathophysiology of schizophrenia, including those related to mitochondrial function. These results suggest that cluster-specific DEGs may underlie some cell type-specific neurobiological changes associated with schizophrenia neuropathophysiology.

### Canonical Pathways

To identify metabolic and cell signaling pathways that are likely to be altered in the primary neuronal clusters, canonical pathway analysis was performed with Ingenuity Pathway Analysis (IPA) using the cluster-specific DEGs. Shared and unique canonical pathways were identified for the primary neuronal clusters (Supplementary table 11). The top five shared canonical pathways for the primary neuronal clusters, as determined by p-value, are presented in Fig. 3a. While the primary neuronal clusters shared predicted disruptions of pathways, the DEGs underlying the effect for each cellular cluster were a combination of shared and unique DEGs (Fig. 3b and Supplementary table 11). These results suggest that a combination of shared and cluster-specific transcriptome alterations drive overall pathway dysfunction.

**Fig. 3:**
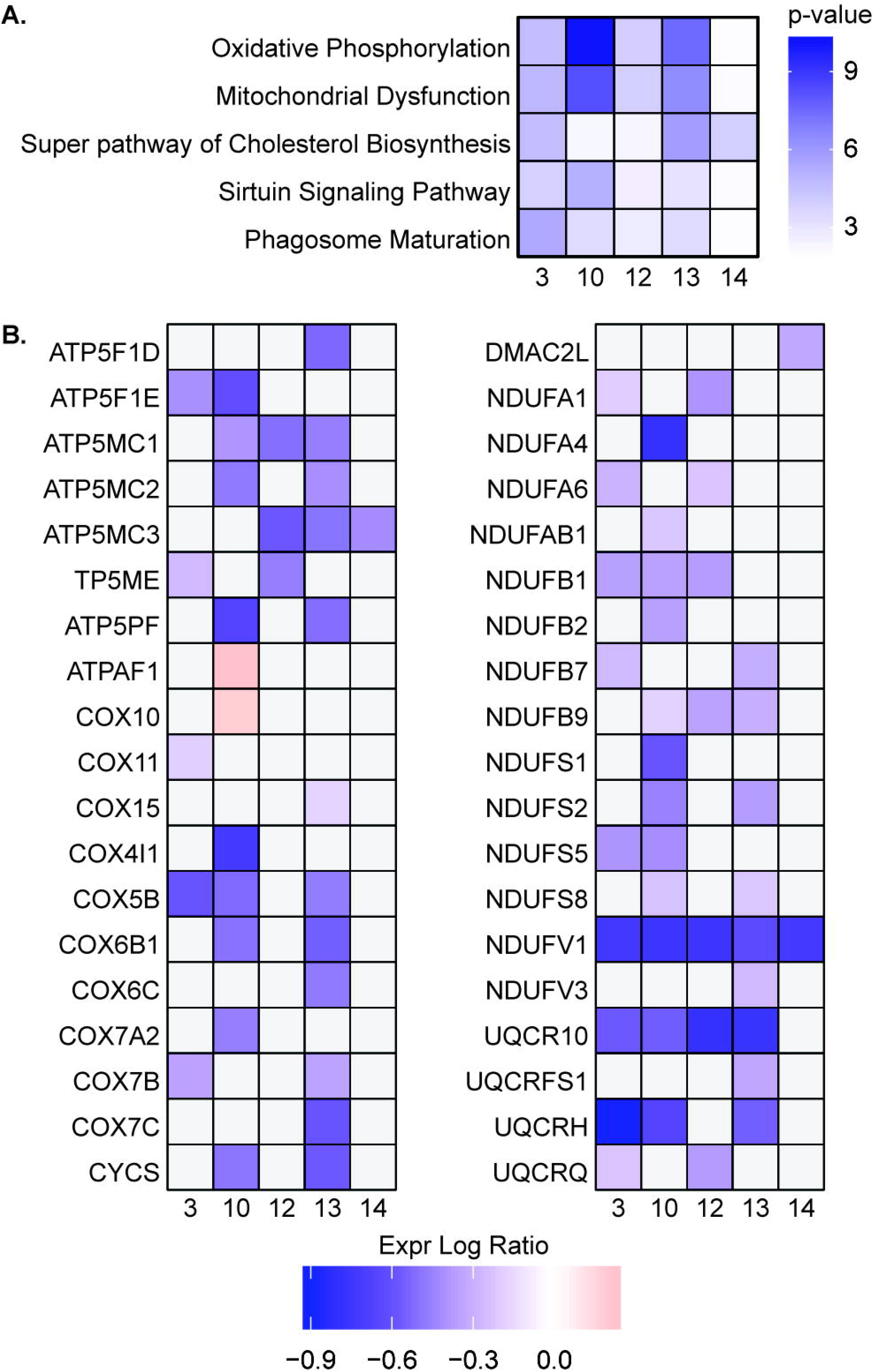
Canonical pathways. **a** Overrepresentation of cluster-specific DEGs in the top five shared canonical pathways for the primary neuronal clusters. **b** Shared and unique cluster-specific DEGs that underlie the oxidative phosphorylation canonical signaling pathway.

### Upstream transcription factors, microRNAs, and regulatory networks

To identify putative mechanisms for the alterations in gene expression observed in the primary neuronal clusters, we assessed overrepresentation of transcription factor (Supplementary table 12) and microRNA (Supplementary table 13) targets among cluster-specific DEGs. Unique transcription factor and microRNA targets were identified for each of the primary neuronal clusters, suggesting potential cell type-specific mechanism for transcriptional dysregulation. Additionally, a subset of transcription factors and microRNAs had significantly overrepresented targets across the primary neuronal clusters, suggesting a potential shared etiology. Unique and shared putative upstream regulatory networks were also identified for all clusters (Supplementary table 14). The top five shared upstream network master regulators for the primary neuronal clusters, as determined by p-value, are presented in Fig. 4a. These predictions were driven by shared and cluster-specific DEGs (Fig. 4b). These results are supported by the direct detection of predicted alterations for a portion of the upstream regulators. For example, *DDX5* function was predicted to be inhibited in four of five primary neuronal clusters (Fig. 4a) and *DDX5* was found to be significantly downregulated in both cluster 3 PVALB+ InNeuro and cluster 13 layer 4/5 ExNeuro (Supplementary table 14).

**Fig. 4:**
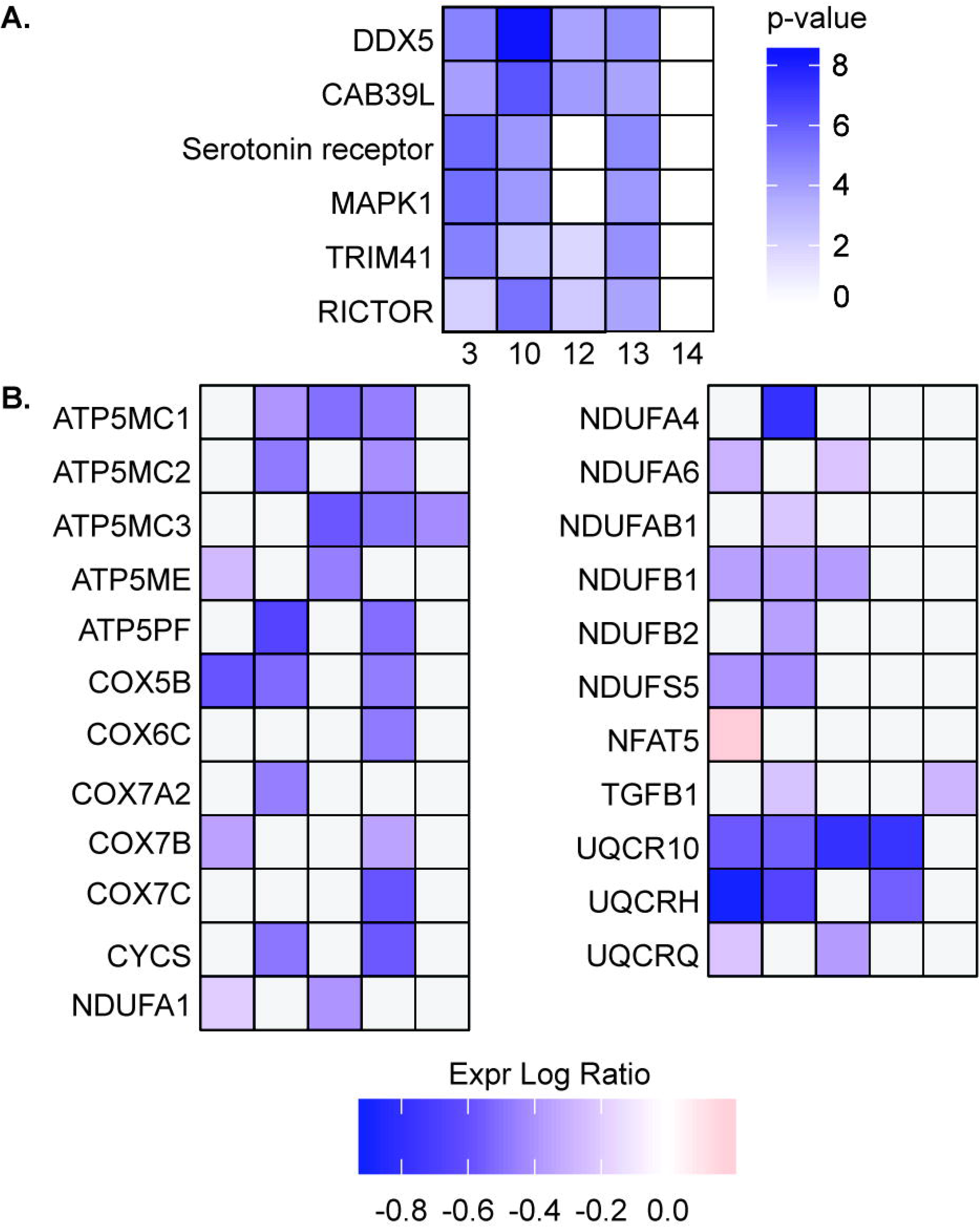
Upstream regulators. **a** Overrepresentation of cluster-specific DEGs in the top five predicted upstream master regulators. **b** Shared and unique cluster-specific DEGs that underlie the prediction of DDX5 as an upstream regulator of cell type-specific DEGs.

## Discussion

This study examined the transcriptome of ~275,000 single nuclei from the dlPFC of persons with schizophrenia and controls. The median UMI and gene counts per nucleus were approximately double those of three recent human postmortem cortical snRNAseq studies, although the ratio of median UMI count to median gene count was similar^14–16^. Differences between versions 2.0 and 3.0 of the 10x Genomics gene expression assay explain at least a portion of our increased gene and UMI yield. The identification of 20 transcriptomically distinct cellular clusters is consistent with other snRNAseq studies of human postmortem cortex^13, 14, 16^ and the number of clusters per major cell type closely approximates those of a recent studies of dlPFC, including identification of a single *HTR2C*+ cluster of pyramidal ExNeuro and two *PVALB*+ clusters of InNeuro^16^. 4,766 differential expression events were detected in 16 of 20 cellular clusters, with ~96% of DEGs occurring in five neuronal cell types. Prior evidence suggests that GWAS loci and gene sets associated with schizophrenia are primarily expressed in a limited subset of neurons, including PVALB+ InNeuro and glutamatergic pyramidal neurons^31^. Similarly, GWAS loci-related genes were overrepresented in the DEGs of cluster 3 *PVALB*+ InNeuro and the cluster 10 layer 5 *HTR2C*+, cluster 12 layer 5, and cluster 14 layer 2/3 ExNeuro. Cluster 12 layer 5 ExNeuro also had an overrepresentation of bipolar disorder GWAS loci, fitting prior knowledge that schizophrenia and bipolar disorder share genetic risk loci. These results support the hypothesis that common genetic variants associated with schizophrenia are relevant in specific sets of neuronal cell types and schizophrenia-related transcriptomic alterations are primarily limited to these cells. Several reports provide fairly consistent evidence of presynaptic marker decreases for frontal cortical fast-spiking parvalbumin +- GABAergic interneurons, coupled with increased postsynaptic GABAA receptors, both of which may be consistent with partial loss of GABAergic inhibition of glutamatergic pyramidal neurons^32, 33^. The large number of DEGs in both cell types provides support for dysfunction of a frontal cortex GABAergic-glutamatergic circuit. No DEG occurred in more than five cellular clusters and that no DEG was present in all the clusters of any multi-clustered major cell type underscores the importance of utilizing single nuclei/single cell approaches to neural transcriptomics.

The substantial overrepresentation of cluster-specific DEGs during critical neurodevelopmental timepoints, 8 to 13 post-conception weeks and 2 years of age (Fig. 2b) supports hypotheses about schizophrenia as a neurodevelopmental disorder ^34–36^. Our analyses identified a substantial number of GO terms and KEGG and canonical pathways related to energy metabolism and oxidative stress in the primary neuronal clusters and prior works have hypothesized the prenatal and early developmental dysregulation of oxidative stress may play a role in the development of schizophrenia, particularly in PVALB+ neurons (reviewed^37^). Taken together, these data suggest potential windows for PVALB+ InNeuro oxidative stress targeted interventions.

Human postmortem studies are identifying increasing numbers of shared and brain region-specific differentially expressed microRNAs that are associated with schizophrenia^38^, including a global increase in microRNA levels^39^. While the experimental approach of this study was unable to directly detect alterations in microRNA expression, complementary approaches to identifying cell type-specific microRNA target overrepresentation yield overlapping predictions of microRNAs known to regulate brain function. For example, miR-424 (aka miR-322) targets were predicted to be overrepresented in the cluster 3 PVALB+ InNeuro and cluster 10 layer 5 HTR2C+ and cluster 13 layer 4/5 ExNeuro. miR-424 is known to regulate BDNF expression^40^ and literature evidence supports a role for alterations in BDNF expression in schizophrenia pathogenesis^41^. Taken together, these data suggest that the neuronal cell type-specific microRNA identified in this study may warrant further investigation.

Several limitations of this study must be noted. First, the relatively small number of postmortem samples analyzed increases the possibility that the subjects are not representative of the broader populations. Replication in a larger sample, including female samples, will be essential for these results. Second, schizophrenia patients frequently have comorbidities (e.g. smoking, obesity) that are less common in control individuals, presenting analytical confounds. Similarly, schizophrenia patients usually have a history of chronic antipsychotic treatment, whereas controls do not. Thus, it is impossible to know at present whether any of the identified DEGs reflect causality or response to chronic pharmacotherapy. It may also be possible to address this issue by studying postmortem brains of persons with schizophrenia who never had antipsychotic treatment. However, at least in the United States, these patients are uncommon. Third, transcriptome-based methods such as snRNAseq have the potential to miss relevant genes that are regulated primarily at the level of translation or splicing, which may also help to shape transcriptomic architecture and be relevant to schizophrenia pathology. Finally, this project was limited to a single brain region from individuals over 18 years of age. Therefore, spatial and temporal changes in gene expression occurring over the course of the disease would not be identified in our analysis. The findings from this study of dlPFC cannot be extrapolated to other areas of the brain, justifying the need for more comprehensive studies. There is also substantial evidence for neurodevelopmental origins in schizophrenia^42, 43^, suggesting there are relevant transcriptomic differences between cases and controls before adulthood. In summary, we have begun to characterize transcriptome alterations in schizophrenia at the level of single neural cells and extension of this work may provide a new basis for the development of effective treatment strategies.

## Availability of Data

Raw sequencing data and sample annotations are available at NCBI GEO accession # GSE158516.

## Acknowledgements

BCR is supported, in part, by a 2017 NARSAD Young Investigator Grant (#26634) from the Brain and Behavior Research Foundation as the Patrick A. Coffer Investigator, funding for which was generously provided by Ronald and Kathy Chandonais. BCR and AEW were supported during part of this work by T32MH014654 (PI is WHB). This work was funded by a National Institutes of Health grant to WHB (R01MH109260). LMS was supported by NIDDK F32 DK118818. MRH was supported by NIDDK R01 DK115762. The authors thank Rachel Kember for her assistance with R. The authors would also like to thank Jane Gaisinsky for her help with tissue preparation. Additionally, the authors thank all the participants, and their families, who donated their brain tissue to research, for making these studies possible.

## Author Contributions

BCR conceptualized this study and performed experiments. BCR and RCC performed bioinformatics and wrote the manuscript. GA contributed to the performance of experiments and manuscript preparation. LMS, AEW, GAD, TNF and MRH contributed to data interpretation and manuscript preparation. WHB provided oversight and contributed to data interpretation and manuscript preparation.

