## Supplemental for "Single-nuclei transcriptomics of schizophrenia prefrontal cortex primarily implicates neuronal subtypes"

**Supplementary Figures**

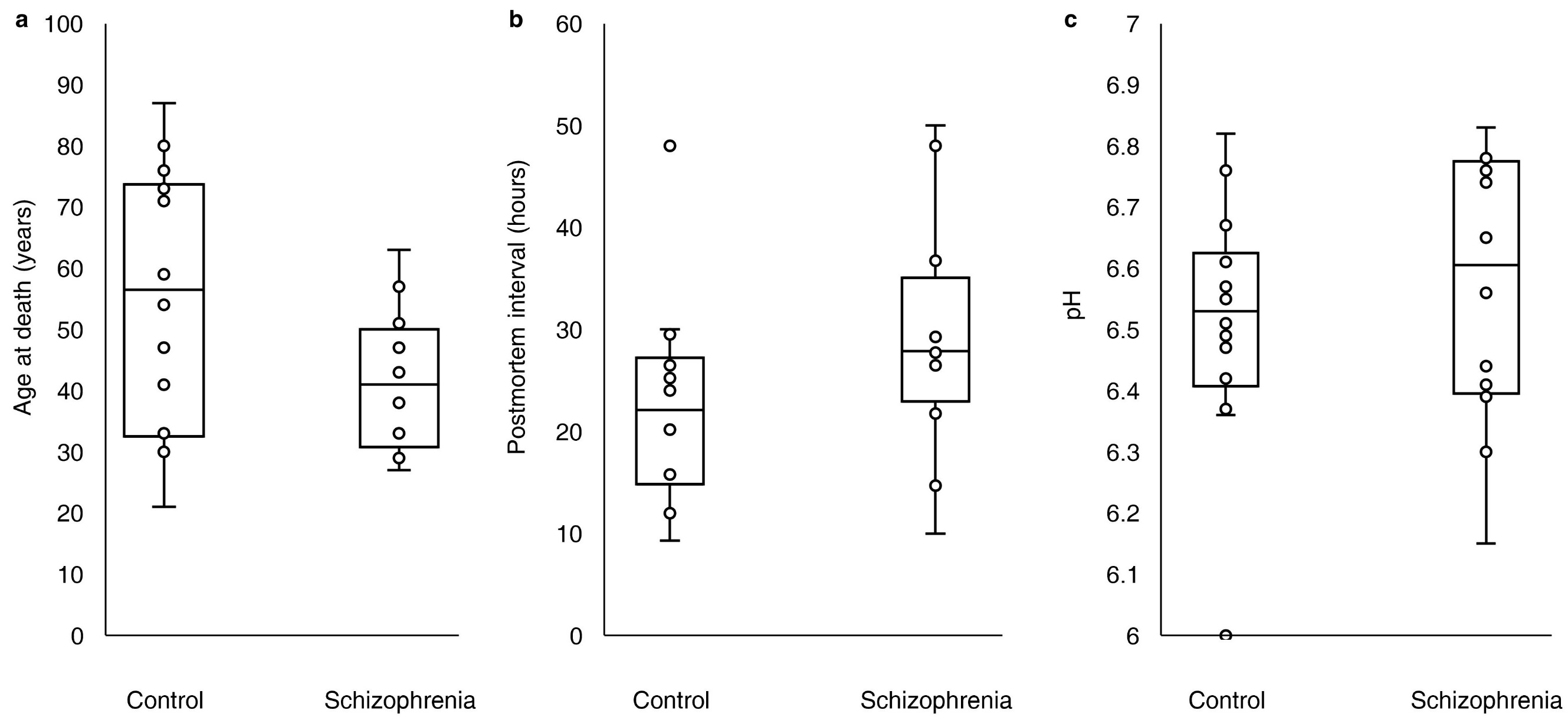

**Supplementary Figure 1: Postmortem sample characteristics**. This study utilized human postmortem brain samples from 12 individuals with schizophrenia and 14 controls. Brain samples for the schizophrenia and control groups did not differ in (a) age at time of death (control 54.4 ± 20.4; schizophrenia 41.7 ± 10.9; Mann-Whitney U = 52.5, z = 1.59, p = 0.11184) , (b) postmortem interval (PMI; control 22.0 ± 9.7; schizophrenia 29.1 ± 11.2; Mann-Whitney U = 50.0, z = -1.72, p = 0.08544), or (c) prefrontal cortex pH (control 6.5 ± 0.2; schizophrenia 6.6 ± 0.2; Mann-Whitney U = 72.5, z = -0.57, p = 0.56868). Data presented as Mean ± Standard Deviation. Box and Whisker plots display the median (center line), first and third quartiles (box), and the maximum and minimum values within 1.5x the interquartile distance (whiskers). Circles denote data points in addition to maximum and minimum whiskers.

**Supplementary Figure 2: Sequencing statistics**. snRNAseq was performed on postmortem brain samples from the schizophrenia and control individuals using a the 10x Genomics Chromium platform. Sequencing data was initially analyzed using the 10x Genomics Cell Ranger pipeline (see Methods). (a) The number of sequencing reads per individual did not vary between the schizophrenia and control groups (control 499,276,080.1 ± 97,471,679.6; schizophrenia 484,510,877.8 ± 101,300,698.3; Mann-Whitney U = 83, z = -0.03, p = 0.97606). (b) The number of nuclei identified per sample (control 13,654.4 ± 2,980.2; schizophrenia 14,210.0 ± 2,249.9; Mann-Whitney U = 76, z = -0.39, p = 0.69654) and (c) the average number of sequencing reads per nuclei per sample (control 37,924.9 ± 10,060.5; schizophrenia 34,501.4 ± 7,189.6; Mann-Whitney U = 70.5, z = 0.67, p = 0.50286) did not differ between the schizophrenia and control groups. Nuclei from the schizophrenia and control groups did not differ in (d) the median number of genes detected per nuclei (control 3,126.8 ± 835.2; schizophrenia 2,825.9 ± 702.3; Mann-Whitney U = 64, z = 1.00, p = 0.31732) or (e) the median number of UMI per nuclei (control 7,301.4 ± 2,853.5; schizophrenia 6,403.0 ± 2,408.0; Mann-Whitney U = 72, z = 0.59, p = 0.5552). Data presented as Mean ± Standard Deviation. Box and Whisker plots display the median (center line), first and third quartiles (box), and the maximum and minimum values within 1.5x the interquartile distance (whiskers). Circles denote data points in addition to maximum and minimum whiskers.

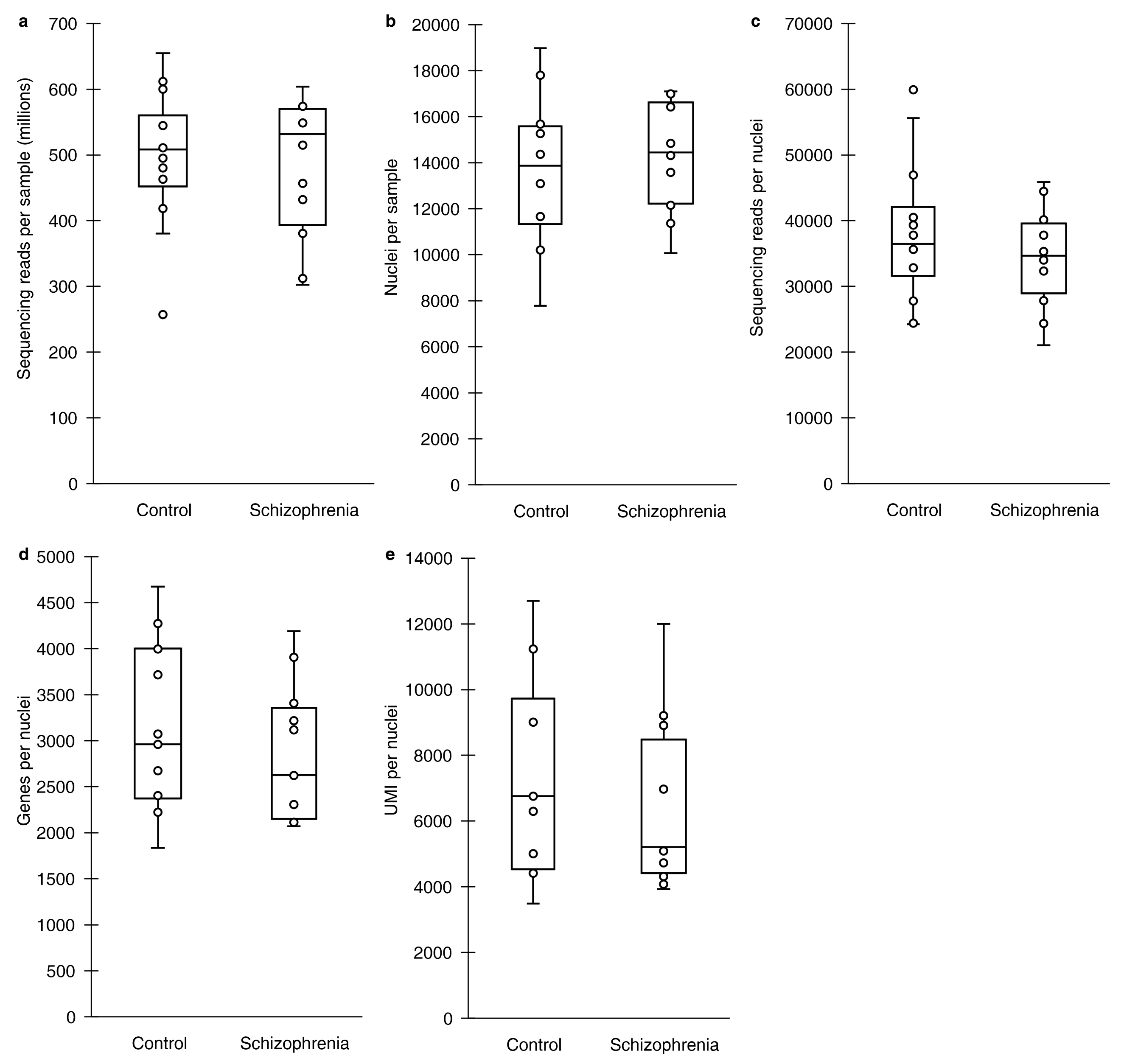

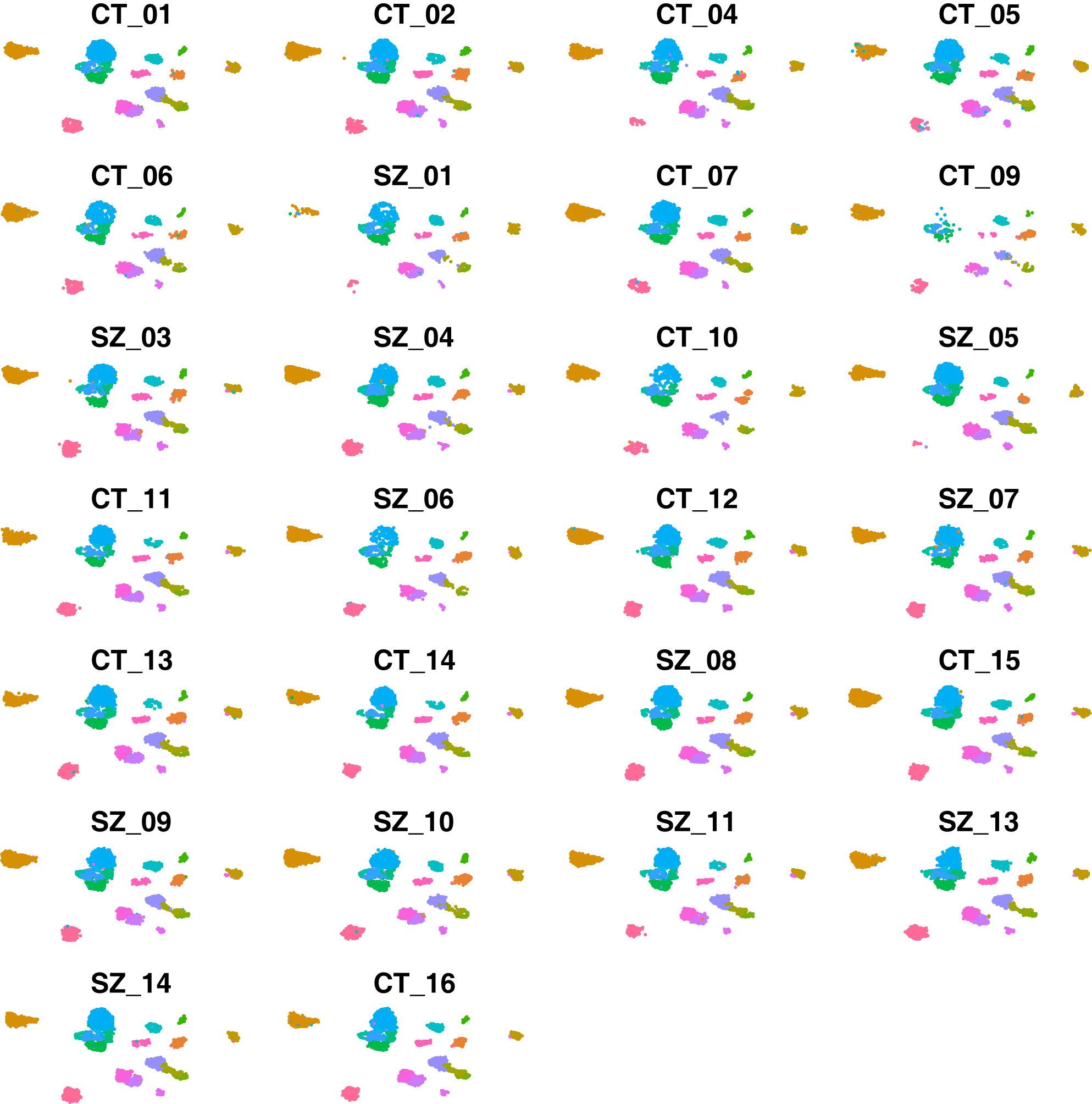

**Supplementary Figure 3: Individual UMAP**. UMAP plots for all individuals showing their contribution to the clusters.

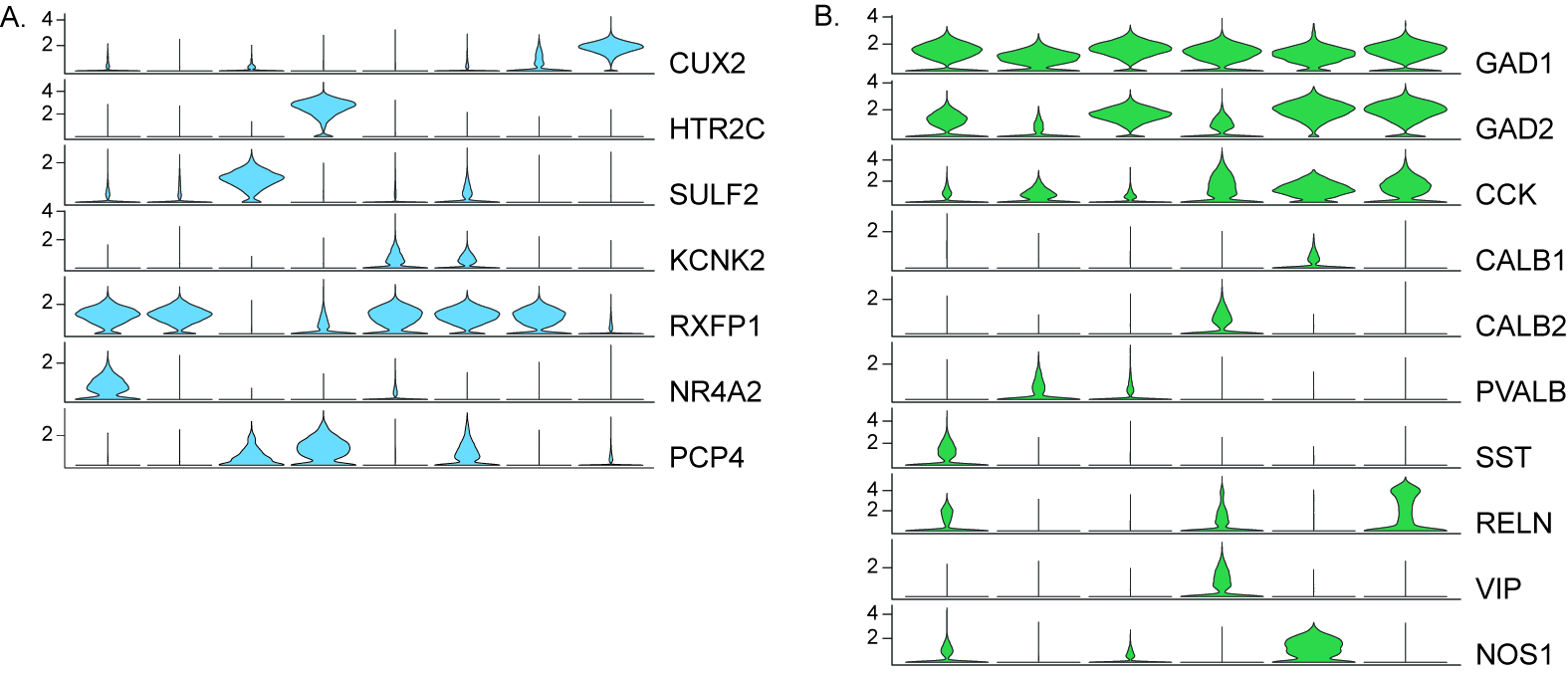

**Supplementary Figure 4: Differential expression events.** (a) Excitatory neuron clusters were annotated with known markers of cortical layer II/III (CUX2), layer V (HTR2C, SULF2, KCNK2 and PCP4), layer V/VI (RXFP1), and layer VI (NR4A2) to determine the cortical layer represented by each excitatory neuron cluster. (b) Inhibitory neuron clusters were annotated using known markers of inhibitory neuron subtypes, including calcium binding proteins (PVALB, CALB1, and CALB2), neuropeptides (SST, VIP, and CCK), and other known markers of cortical inhibitory neurons (GAD1, GAD2, RELN, and NOS1).

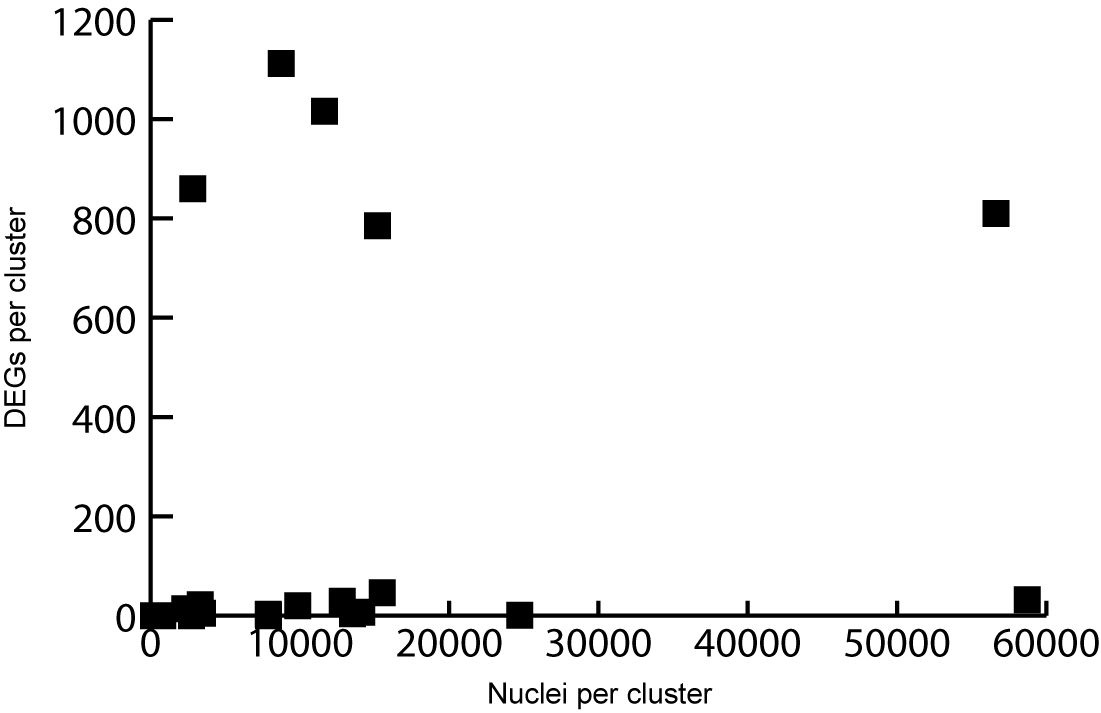

**Supplementary Figure 5:** A scatter plot depicting the lack of relationship between the number of differentially expressed genes (DEGs) detected in a cluster and the number of nuclei present in the cluster (r(18) = 0.16, p = 0.50, Pearson’s Correlation).

**Supplemental Methods**

Isolation of Nuclei

Tissue was dounce homogenized in 1 mL lysis buffer (Nuclease-free water with 10 mM Tris-HCL, 10 mM NaCl, 3 mM MgCl_2_, 0.5% NP-40) using 20 strokes of the loose pestle and 15 strokes of the tight pestle. The homogenate was diluted to 5 mL with lysis buffer and left to lyse on ice for an additional 5 minutes, followed by addition of an equal volume of wash buffer (1X PBS with 2% BSA, 1:1000 RNase inhibitor, 0.25% Glycerol). Samples were then passed through a 30 μm cell strainer and centrifuged at 500 x g for 5 minutes at 4°C. Supernatants were decanted and pellets were resuspended in 10 mL wash buffer. Samples were passed through a 30 μm cell strainer for a second time and centrifuged at 500 x g for 7 minutes at 4°C. Supernatants were decanted and pellets were resuspended in 5 mL wash buffer and centrifuged at 500 x g for 10 minutes at 4°C. Supernatants were decanted, pellets were resuspended in 500 µL wash buffer, and mixed with 500 µL of 50% iodixanol solution (Optiprep), to make a 25% iodixanol solution containing the washed nuclei. The 1 mL 25% iodixanol nuclei solution was layered on 1 mL of a 29% iodixanol cushion and centrifuged at 10,000 x g for 30 minutes at 4°C to pellet nuclei.

Annotation of Cell Clusters

Major cell types were identified using known cell type markers and methods described in Nagy et al.^1^: Microglia: - *SP1, MRC1, CX3CR1*; Endothelial – *CLDN5*; Astrocytes – *GLUL, SOX9, AQP4, GJA1, NDRG2, GFAP, ALDH1A1, ALDH1L1*; Oligodendrocyte Precursor Cells – *PDGFRA, PCDH15, OLIG1, OLIG2*; Oligodendrocytes – *PLP1, MAG, MOG, MOBP, MBP*; Excitatory Neurons – *SATB2, SLC17A7*; Neurons - *SNAP25, STMN2, RBFOX3*; Inhibitory Neurons – *GAD1, GAD2, SLC32A1*. The cortical layer of excitatory neuron clusters was identified using previously described layer markers^1^: Layer II: *GLRA3*; Layer II/III: *CUX2, CARTPT, LAMP5*; Layer V: *KCNK2, SULF2, PCP4, HTR2C, FEZF2*; Layer V/VI: *RXFP1, TOX, ETV1, RPRM, FOXP2*; Layer VI: *NR4A2, SYNPR, NTNG2*. Similarly, inhibitory neuron cell types were identified using previously described markers^1^: *GAD1, GAD2, CCK, CALB1, CALB2, PVALB, SST, RELN, VIP, NOS1*.

1.	Nagy C, Maitra M, Tanti A, Suderman M, Theroux JF, Davoli MA *et al.* Single-nucleus transcriptomics of the prefrontal cortex in major depressive disorder implicates oligodendrocyte precursor cells and excitatory neurons. *Nat Neurosci* 2020; **23**(6)**:** 771-781.
